# OspA Antibodies Inhibit the In vitro Transmigration of *Borreliella burgdorferi*

**DOI:** 10.1101/2025.11.20.689610

**Authors:** Ankita Bhattacharyya, Nicholas J. Mantis

## Abstract

Antibodies against Outer surface protein A (OspA) block tick-to-mammal transmission of the Lyme disease spirochete, *Borreliella burgdorferi*. Evidence suggests that antibodies ingested by a tick during a blood meal entrap OspA-expressing *B. burgdorferi* within the tick midgut and prevent spirochete migration to the salivary gland and host dermis. In this study, we employed a recently described Transwell system to quantitate the impact of OspA antibodies on *B. burgdorferi* transmigration. We demonstrate that spirochete movement from the lower Transwell chamber to the upper chamber over a 20-hour period was reduced by ≥99% in the presence of the transmission-blocking mouse monoclonal IgG antibody (mAb) LA-2. In the case of LA-2, transmigration inhibition coincided with the formation of large antibody-induced bacterial aggregates in the lower Transwell chamber, as determined by dark field microscopy and flow cytometry. Further examination of a panel of human OspA mAbs of varying affinities and epitope specificities revealed that virtually all could block *B. burgdorferi* transmigration, although most did so in the absence of measurable spirochete agglutination. We propose that transmigration arrest may represent a possible mechanism by which OspA antibodies entrap *B. burgdorferi* in the tick midgut and limit transmission to mammalian hosts.

## INTRODUCTION

The spirochete *Borreliella burgdorferi* sensu lato is the etiologic agent of Lyme disease, the most prevalent tick-borne disease in the United States (Steere *et al*., 2016). *B. burgdorferi* is 10-20 microns in length (0.3 microns wide) and swims with a planar (flat wave) morphology driven by 7-10 periplasmic flagella (Goldstein *et al*., 1994, Charon & Goldstein, 2002). Motility enables the spirochete to disseminate in a variety of viscous environments and tissues (Harman *et al*., 2012, Harman *et al*., 2013, Bockenstedt *et al*., 2014). Motility is paramount in the context of tick-mediated transmission in that the spirochete migrates during the course of a blood meal from the tick midgut to the hemocoel and salivary glands before being deposited into the dermis of the mammalian host (De Silva & Fikrig, 1995). Breaching the tick midgut epithelium is a particularly dynamic process that occurs over a 24-72 h period, as revealed through in situ confocal imaging of fluorescent spirochetes (Dunham-Ems *et al*., 2009). The importance of motility to *B. burgdorferi*’s infectious life cycle is underscored by the fact that flagella mutants are not only attenuated for mammalian infection but inept for tick to mouse transmission (Sultan *et al*., 2013, Hyde, 2017).

During egress from the tick midgut, *B. burgdorferi* is also vulnerable to host factors, including antibodies. In fact, Lyme disease vaccines and monoclonal antibodies directed against Outer surface protein A (OspA) afford protection by entrapping spirochetes within the tick midgut during a feeding event (Fikrig *et al*., 1990, Fikrig *et al*., 1992, de Silva *et al*., 1996, de Silva *et al*., 1999, Gipson & de Silva, 2005, Srivastava & de Silva, 2008). OspA is a ∼30 kDa lipoprotein expressed at high levels on the surface of *B. burgdorferi* that reside within the midgut of unfed ticks. OspA is downregulated as the spirochete migrates to the salivary gland and eventually extinguished when the bacteria are within mammalian hosts (Srivastava & de Silva, 2008). It was recognized almost three decades ago in mouse (and other rodent) models that circulating OspA polyclonal and monoclonal antibodies (mAbs) are highly effective at blocking tick-mediated transmission of *B. burgdorferi* (Fikrig *et al*., 1990, Schaible *et al*., 1990). These and other observations eventually led to the development of first- and second-generation human and veterinary OspA-based Lyme disease vaccines (Steere *et al*., 1998, Gomes-Solecki *et al*., 2020, Wormser, 2022).

The molecular mechanisms by which OspA antibodies interfere with *B. burgdorferi* transmission are not fully understood, but evidence suggests that the interactions likely occur within the midgut and are not dependent on human complement (de Silva *et al*., 1996, de Silva *et al*., 1999, Pal *et al*., 2000, Rathinavelu *et al*., 2003, Gipson & de Silva, 2005). Since those early reports very little has been done in this area, probably because of the technical challenges associated with sorting out antibody-spirochete interactions within the context of tick tissues (Dunham-Ems *et al*., 2009). However, Van Gundy and colleagues recently developed a Transwell method to evaluate *B. burgdorferi* migration in response to chemotactic gradients, including salivary proteins (Van Gundy *et al*., 2021). Transwell units consist of an insert with a microporous membrane (3.0 micron) that sits within 96-well microtiter plates.

Spirochetes suspended in a low-nutrient medium and seeded in the lower chamber of 96 well Transwell units migrate against gravity into the upper chamber in response to chemoattractant, including BSK-II complete medium (Van Gundy *et al*., 2021). In this report, we employed the Transwell assay as a model system to investigate the impact of OspA antibodies on *B. burgdorferi* migration across a physical barrier.

## MATERIALS AND METHODS

### Strains and culture conditions

*B. burgdorferi* strain B31-A, a high-passage, non-infectious clone of wild-type B31 carrying the plasmid pBSV2G_P*_flaB_*-msfGFP (Addgene 118231) that constitutively expresses GFP expression, was used for most of the assays (Burgdorfer *et al*., 1982, Takacs *et al*., 2018). For the assays with different OspA serotypes, *B. burgdorferi* strain HB19-R1, a high-passage derivative of human blood isolate HB19 lacking linear plasmid 54 (Δ*ospAB*) was transformed with the IPTG-inducible mScarlet-I reporter plasmid expressing each of the seven different OspA variants (ST 1-7), regulated by the P*_ospAB_* promoter from *B. burgdorferi* B31 (Willsey et al, *manuscript submitted*) (Sădziene *et al*., 1992). All strains were maintained as frozen glycerol stocks at - 80°C and cultured routinely in gelatin-free BSK-II complete media, supplemented with 50 µg/mL of gentamicin, in microaerophilic and static growth conditions at 33°C with 2% CO_2_, unless otherwise stated.

### Monoclonal antibodies (mAbs)

The OspA-specific mAbs used in this study (**Table 1**) were expressed and purified by Dr. Lisa Cavacini (UMass Chan Medical School, Worcester, MA), as described (Haque *et al*., 2022). Chimeric LA-2 IgG1 was custom synthesized by ZabBio, Inc (San Diego, CA).

**Table 1:** Functional characteristics of mAbs used in this study.

| **Table 1: Functional characteristics of mAbs used in this study** |  |  |  |  |
| --- | --- | --- | --- | --- |
| **Bin** | **mAb<sup>a</sup>** | **IEDB ID<sup>b</sup>** | **migration inhibition (%)<sup>c</sup>** | **agglutination (%)<sup>c</sup>** |
| 1 | <u>857-2</u> | 2144660 | 46 ± 5 | 4 ± 0.8 |
| 1 | <u>221-7</u> | 1597628 | 62 ± 13 | 4 ± 2.2 |
| 1 | 221-5 | pending | 55 ± 7 | 16 ± 5 |
| 1 | 221-11 | 2150407 | 61 ± 9 | 2 ± 0.4 |
| 1 | 221-20 | 2150407 | 37 ± 25 | 23 ± 11 |
| 1 | 223-5 | pending | 50 ± 10 | 1 ± 0.2 |
| 1 | 227-2 | 2150577 | 70 ± 3 | 2 ± 0.3 |
| 2 | <u>212-55</u> | 2151022 | 67 ± 10 | 3 ± 2 |
| 2 | 319-28 | 2151021 | 33 ± 12 | 2 ± 0.4 |
| 3 | <u>LA-2</u> | na | 98 ± 1 | 15 ± 6 |
| 3 | <u>319-44</u> | 2144728 | 89 ± 4 | 6 ± 4 |
| 3 | 319-33 | 2150778 | 83 ± 3 | 34 ± 10 |
| 3 | 3-24 | 2151024 | 93 ± 4 | 32 ± 11 |

<sup>a</sup>, Underline indicates mAbs with transmission blocking activity in mouse model; <sup>b</sup>, IEDB ID, https://www.iedb.org; <sup>c</sup> mean + SD of three independent replicates.

**Table 2:** V_H_Hs used in this study.

| Table 2: V<sub>H</sub>Hs used in this study |  |  |  |
| --- | --- | --- | --- |
| Bin | V<sub>H</sub>H/VHH-Fc | IEDB ID <sup>a</sup> | Epitope |
| 1 | L8H8 | 2257224 | β-Strands 10-14 |
| 1 | L8D3 | 2257226 | β-Strands 11-13 |
| 1 | L8G12 | 2257227 | β-Strands 11-14 |
| 3 | L8A9 | pending | β-Strands 16-18 |
| 3 | L8C3 | pending | β-Strands 16-18 |
| 3 | L8D9 | pending | β-Strands 16-18 |
| <sup>a</sup>. IEDB ID, https://www.iedb.org |  |  |  |

### Transmigration assays

The Transwell^®^ migration assays were performed essentially as described by Van Gundy (Van Gundy *et al*., 2021), with several modifications. HTS Transwell^®^ 96 well permeable supports (Corning, Corning, NY) with tray inserts containing 3.0 µm pore size polycarbonate membrane in each well were used for the assays. Frozen glycerol stocks of respective *B. burgdorferi* strains were thawed and cultured for ∼3 days, then diluted in fresh medium at a starting cell concentration of 1.5×10^6^ spirochetes/mL and grown till mid-exponential phase (2-5×10^7^ spirochetes/mL; ∼40-42 h). Spirochetes were collected by centrifugation at 1800×*g* at room temperature. The cell pellet was washed once with phosphate buffered saline (PBS), pH 7.4, and then resuspended in the low-nutrient, methylcellulose-free *Borrelia* motility buffer (Zhang & Li, 2018), supplemented with 50 µg/mL of gentamicin. For the HB19-R1 strains, the buffer was also supplemented with 1 mM IPTG for inducing mScarlet production.

Cell suspensions were added to the lower chambers (LC) of the Transwell units along with appropriate dilutions of mAbs (**Table 1**) or polyclonal serum that had been heat-inactivated at 56°C for 30 min. In the end, the lower chamber contained 2×10^6^ HB19-R1 spirochetes or 2×10^7^ B31-A spirochetes in 200 µL. The upper chambers (UC) of the Transwell unit were filled with 150 µL of BSK-II complete medium, free of gelatin and Phenol Red, and supplemented with 50 µg/mL of gentamicin (and 1 mM IPTG for the HB19-R1 strains). The Transwell plate was sealed with Parafilm^®^ and incubated at 33°C with 5% CO_2_ for 18-20 h to enable spirochete migration across the polycarbonate membrane from the lower to the upper chamber. After the incubation, 75 µL volume was collected, without touching the membrane, from the upper chamber of each well, and diluted in 425 µL of PBS (total volume 0.5 mL) in a Falcon^®^ 5 mL polystyrene test tube (Corning) for flow cytometry. The transmigration assays were done with 3-5 technical replicates (wells/plate) per assay and the assays were independently repeated three times. Technical replicates were averaged for each independent assay and average migration of spirochetes in antibody treated conditions were normalized to that of untreated controls for statistical analyses (Blainey *et al*., 2014).

### Flow cytometry

To quantitate migration, B31-A spirochetes were enumerated using a FACSCalibur (BD Biosciences) for 1 min at the ‘Lo’ flow rate setting (12 µL/min), with an excitation wavelength of 488 nm and a 530/30 emission filter (FL1). Events were gated to exclude debris, based on their size (forward scatter, FSC), granularity (side scatter, SSC), and GFP fluorescence intensity, using the BD CellQuest™ Pro software. HB19-R1 spirochetes, on the other hand, were counted on a BD FACSymphony A3 for 1 min at the ‘Hi’ flow rate setting (60 µL/min) with an excitation wavelength of 561 nm and a 610/20 emission filter (PE-CF594). Events were gated based on FSC, SSC, and mScarlet fluorescence intensity, using the BD FACSDiva™ software. We used time (at a specific flow rate) as the acquisition parameter for measuring migration to maintain uniformity between control and treated samples. We optimized the time to 1 min as it was sufficient to collect a measurable number of fluorescence events and at a rate that enabled us to run up to 96 samples in succession. Migration, henceforth, is represented as the number of gated GFP or mScarlet events collected over a minute.

To quantitate antibody-mediated spirochete agglutination, aliquots (200 µL) from the lower chamber were added to 300 µL of PBS (total volume 0.5 mL) in a 5 mL polystyrene test tube. A total of 20,000 events were collected by gating to exclude debris, and size and granularity of events were measured using FSC-SSC dot plots with appropriate quadrants, as described (Frye *et al*., 2022). Agglutination was represented as the percentage of events having increased FSC and SSC due to aggregation (sum of events in the lower-right, upper-left, and upper right quadrants) relative to free cells (events in lower-left quadrant). Percent agglutination of all treatments was normalized relative to untreated controls.

Analysis of normalized data was done using GraphPad Prism version 10.6.0 for Windows (GraphPad Software, Boston, MA). Statistically significant differences between treatments and their respective controls were determined using Ordinary one-way ANOVA followed by Dunnett’s or Tukey’s multiple comparisons *post hoc* test.

### Dark-field Microscopy

For examination of the spirochetes in the lower chamber of the Transwell units, 7 µL aliquots were spotted onto glass microscope slides and covered with cover glass (22×22, No. 1, 0.13-0.16 mm thickness). Spirochetes were imaged using an Olympus BH-2 RFCA upright microscope (Evident Scientific; Waltham, MA) paired with a 20X objective (N.A. 0.70) in a dark-field setting. Images and videos were acquired using an Excelis™ MPX-20RC microscopy camera (Accu-Scope, Commack, NY) paired with CaptaVision+ software version 2.4.9.0 for image analysis.

## RESULTS

Van Gundy and colleagues developed a Transwell^®^ assay to quantify the transmigration of *B. burgdorferi* in response to chemotactic factors, like tick salivary gland proteins (Van Gundy *et al*., 2021). Transwells consist of an insert with a microporous membrane (3.0 micron) that sits within 96-well microtiter plates. Spirochetes suspended in a low-nutrient medium and seeded in the lower chamber of 96 well Transwell units migrate against gravity into the upper chamber in response to chemoattractant, including BSK-II complete medium (Van Gundy *et al*., 2021). We sought to adopt this system to assess the impact of OspA antibodies on spirochete movement across a physical barrier such that might be encountered during transmission (**Figure 1A**).

**Figure 1.**
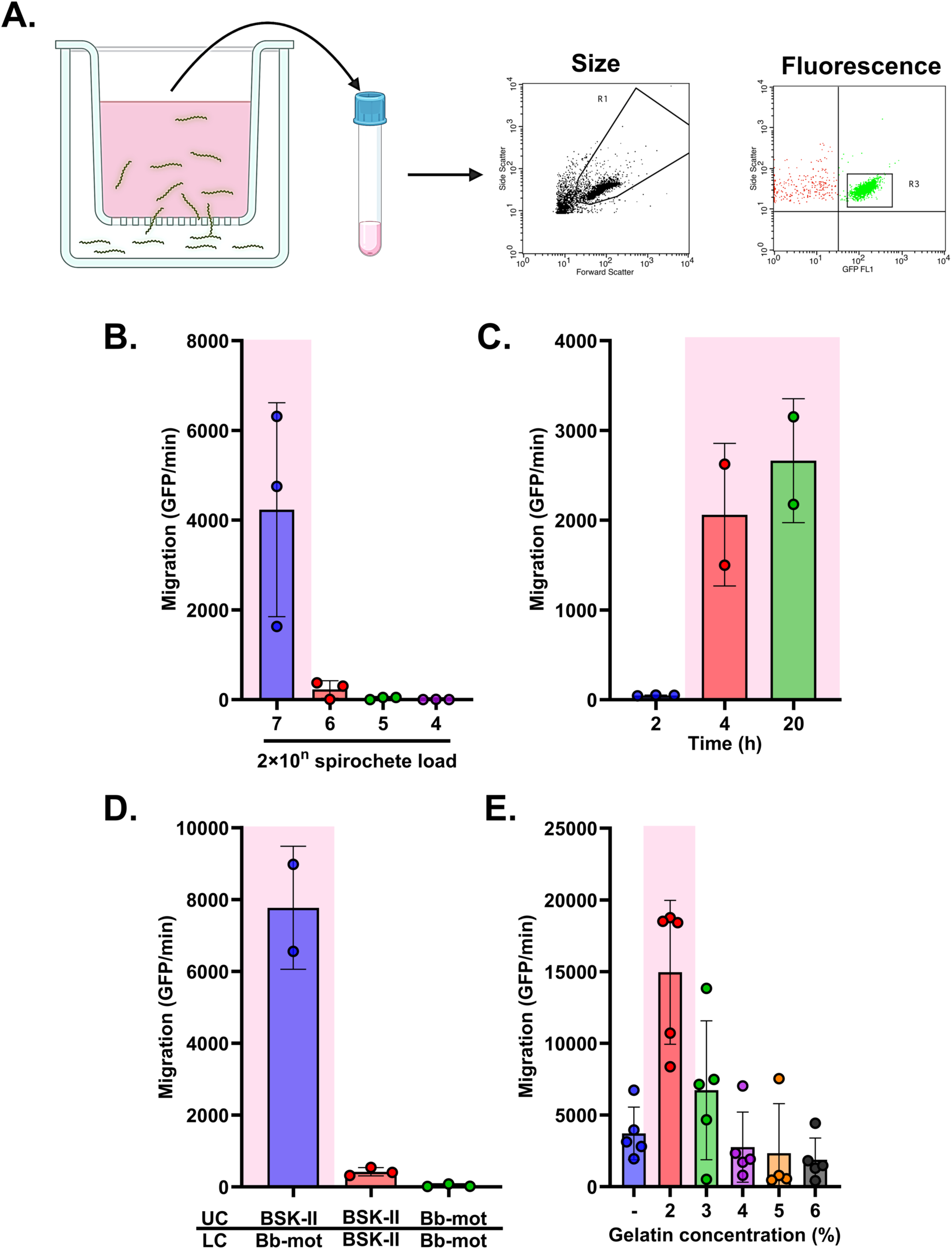
An *in vitro* Transwell system optimized to measure the impacts of OspA antibodies on spirochete migration. **(A)** Fluorescent-labeled spirochetes were suspended in a low-nutrient *Borrelia* motility buffer (Dulbecco’s PBS, 2% recrystallized bovine serum albumin, 0.1 mM EDTA, pH 7.4) and seeded into the Transwell lower chamber and allowed to migrate to the upper chamber containing gelatin- and Phenol Red-free BSK-II media. Migration was measured by collecting upper chamber sample and counting fluorescence events for a minute through flow cytometry using effective gating strategies for size and fluorescence. **(B)** Data represent migration of spirochetes (GFP events/min) relative to a range of spirochete concentrations seeded in the Transwell lower chamber. **(C)** Data represent migration of spirochetes (GFP events/min) relative to a range of Transwell incubation periods. **(D)** Migration of spirochetes (GFP events/min) relative to a chemical gradient. Spirochetes were either suspended in gelatin-free BSK-II media or *Borrelia* motility buffer (Bb mot) and seeded in the lower chamber (LC), with upper chamber (UC) containing either gelatin- and Phenol Red-free BSK-II media or *Borrelia* motility buffer (Bb mot). **(E)** Migration of spirochetes (GFP events/min) in the presence of a range of gelatin concentrations layered over the polycarbonate Transwell membrane, compared to the absence of gelatin. Data in all charts are representative, acquired from a single biological replicate with three technical replicates. Error bars in all charts represent SD of the mean. Statistical significance was determined by one-way ANOVA followed by Tukey’s (B, C, D) or Dunnett’s (E) *post hoc* multiple comparisons test, as applicable. Pink shading indicates *p*≤0.05.

In our model, *B. burgdorferi* B31-A cells expressing GFP or HB19-R1 cells expressing mScarlet were suspended in low-nutrient motility buffer and seeded in the lower chamber of 96 well Transwell units at a range of concentrations (10^4-7^ total) for 2, 4 or 20 h (**Figure 1B, C**). We found that maximal migration was observed when 2×10^7^ spirochetes were seeded in the chamber (**Figure 1B**) and allowed to migrate for 20 h (**Figure 1C**). Migration was dependent on a chemoattractant, as spirochetes were detected in the upper chamber when it contained BSK-II medium but not *Borrelia* motility buffer consisting of only Dulbecco’s PBS, bovine serum albumin,and EDTA. Furthermore, no measurable migration of spirochetes from the lower to the upper chamber was observed when BSK-II or motility buffer were on both sides of the membrane, demonstrating that a chemoattractant gradient is necessary for *B. burgdorferi* migration (**Figure 1D**).

Van Gundy layered the polycarbonate Transwell membrane with 6% gelatin as a diffusion barrier (Kimsey & Spielman, 1990, Harman *et al*., 2012). We similarly investigated the impact of a range of gelatin concentrations (2-6%) on spirochete migration. Consistent with the literature, *B. burgdorferi* migration was significantly enhanced in the presence of 2% gelatin, compared to absence of gelatin, but reduced at higher concentrations (**Figure 1E**) (Harman *et al*., 2012). The addition of 2% gelatin in our migration assays proved to be a confounding factor, however, because the gelatin liquefied at 33°C and affected both pipetting and background values in the flow cytometer. For that reason, we eventually conducted our assays without the addition of gelatin. It should be stated that it is unclear how a chemoattractant diffusion barrier is maintained in the absence of gelatin, considering that 3.0 micron pores would be permeable to stimulants. We speculate that chemoattractant factors in BSK II may be passively adsorbed to the polycarbonate membrane itself, thereby creating a pseudo-chemoattractant gradient sufficient to drive the observed spirochete migration from the lower chamber to the upper chamber in both a time and dose-dependent manner.

### LA-2 blocks Bb migration in a dose-dependent manner

LA-2 is a well-characterized transmission-blocking OspA mAb (Schaible *et al*., 1990, Wang *et al*., 2016). It targets an epitope on OspA’s C-terminus encompassing *β*-strands 16-21 and has potent complement-dependent borreliacidal activity *in vitro* (Ding *et al*., 2000). To evaluate the impact of LA-2 IgG on *B. burgdorferi* B31-A transmigration, spirochetes were mixed with 100 µg mL^-1^ LA-2 or an isotype control just prior to being seeded in the lower chamber of the Transwell unit. The number of spirochetes in the upper chamber was enumerated 20 h later. We found that LA-2 treatment resulted in >99% reduction in spirochete migration relative to the isotype control (**Figure 2A**). We therefore conducted a dose response analysis in which a fixed number of *B. burgdorferi* B31-A cells were mixed with ten-fold serial dilutions of LA-2 IgG (10 - 0.001 µg mL^-1^).

**Figure 2.**
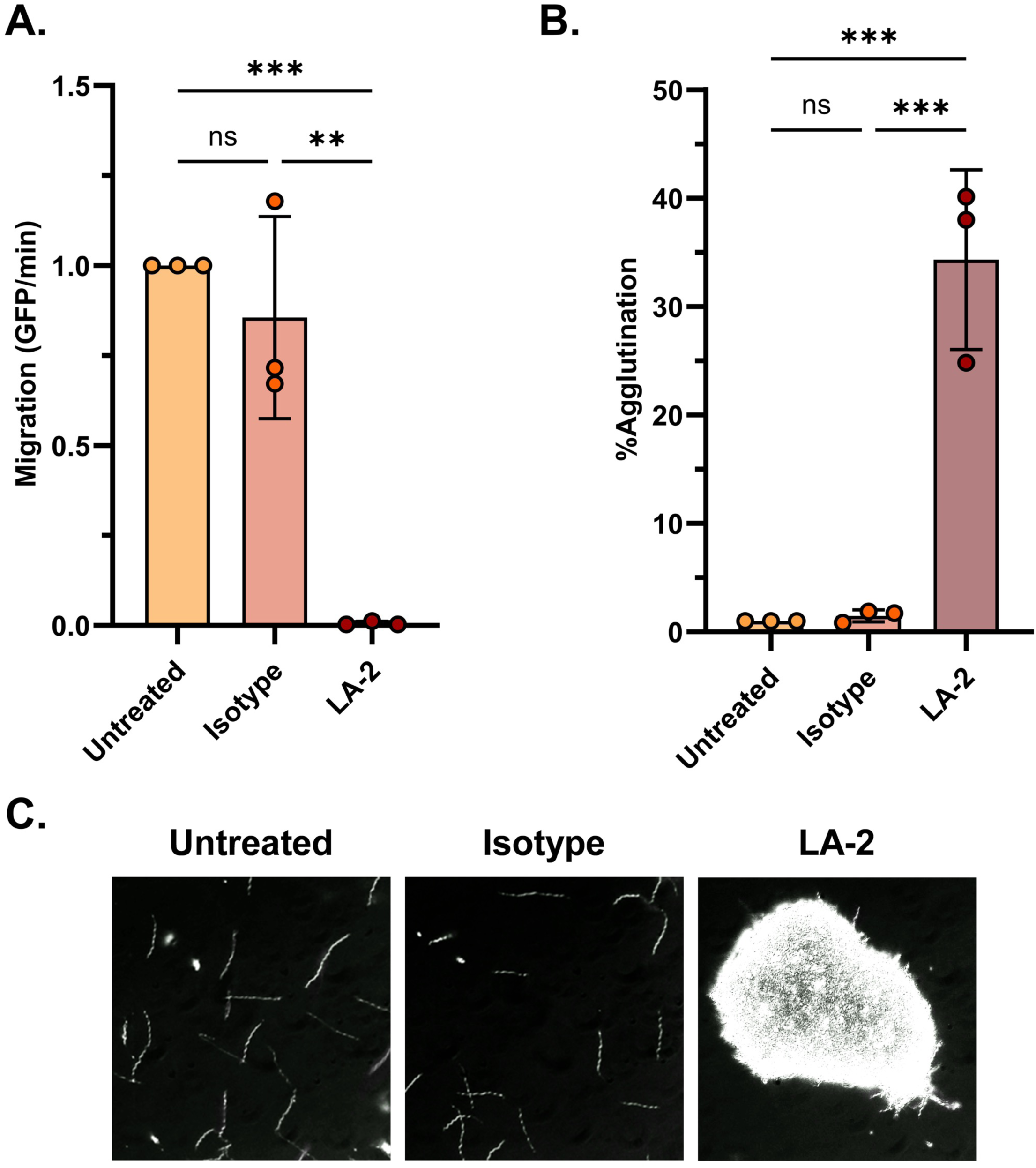
LA-2 significantly inhibits the migration of *B. burgdorferi* B31-A and agglutinates spirochetes in the Transwell assay. 2×10^7^ GFP-tagged spirochetes were either untreated or treated with 100 µg/mL of LA-2 or an isotype control mAb in the Transwell assay (Refer to Materials and Methods). Data were acquired from three independent biological replicates. **(A)** Data represent migration (GFP events/min) normalized to untreated spirochetes and error bars represent standard deviation of the mean. **(B)** Data represent percent agglutination of spirochetes in the lower Transwell chamber normalized to untreated spirochetes and error bars represent standard deviation of the mean. Percent agglutination was calculated by summing increased size (FSC) and granularity (SSC) events relative to the total number of events collected, that represent antibody-mediated aggregation of live spirochetes. Statistical significance was determined by one-way ANOVA followed by Tukey’s *post hoc* multiple comparisons test. Asterisks indicate *p*≤0.05 (**) and *p*≤0.001 (***) and ns=not significant compared to isotype mAb-treated group. **(C)** Dark field microscopy images of spirochetes in the Transwell lower chamber after overnight incubation. All images have been captured at 20X objective magnification.

Within this series, 10 µg mL^-1^ of LA-2 (roughly equivalent to 2×10^6^ molecules per spirochete) resulted in significant inhibition of spirochete migration (∼98%; *p*<0.05), while inhibitory activity diminished at ≤1 µg mL^-1^ (**Figure 3A**). Monovalent LA-2 Fab’ fragments (10 µg mL^-1^) evaluated under identical conditions reduced migration of B31-A by ∼43% relative to the isotype control, although statistical significance was not achieved (**Figure 4A**). From these studies, we conclude that LA-2 is a potent inhibitor of *B. burgdorferi* migration and that maximal effects occur in bivalent form.

**Figure 3.**
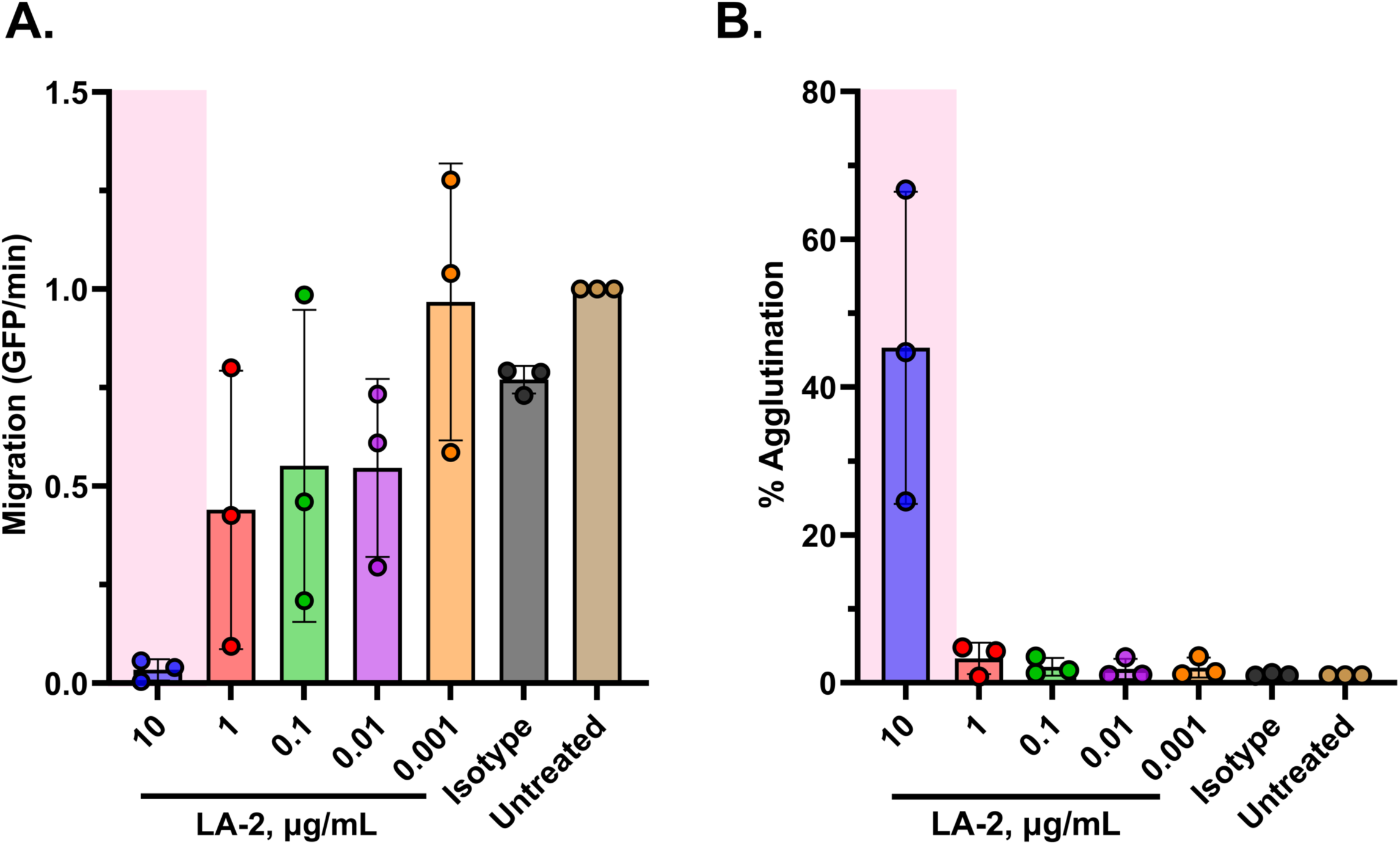
LA-2 inhibits the migration of *B. burgdorferi* B31-A in a dose-dependent manner and migration inhibition correlates with antibody-mediated spirochete agglutination. 2×10^7^ GFP-tagged spirochetes were either untreated or treated with a range of LA-2 concentrations or 10 µg/mL of isotype control mAb in the Transwell lower chamber. The Transwell migration assay was performed as described in Materials and Methods. Data were acquired from three independent biological replicates. **(A)** Data represent migration (GFP events/min) normalized to untreated spirochetes and error bars represent standard deviation of the mean. **(B)** Data represent percent agglutination of spirochetes in the lower Transwell chamber normalized to untreated spirochetes and error bars represent standard deviation of the mean. Percent agglutination was calculated as described in Materials and Methods. Statistical significance was determined by one-way ANOVA followed by Dunnett’s *post hoc* multiple comparisons test. Pink shading indicates *p*≤0.05 compared to isotype mAb-treated group.

**Figure 4.**
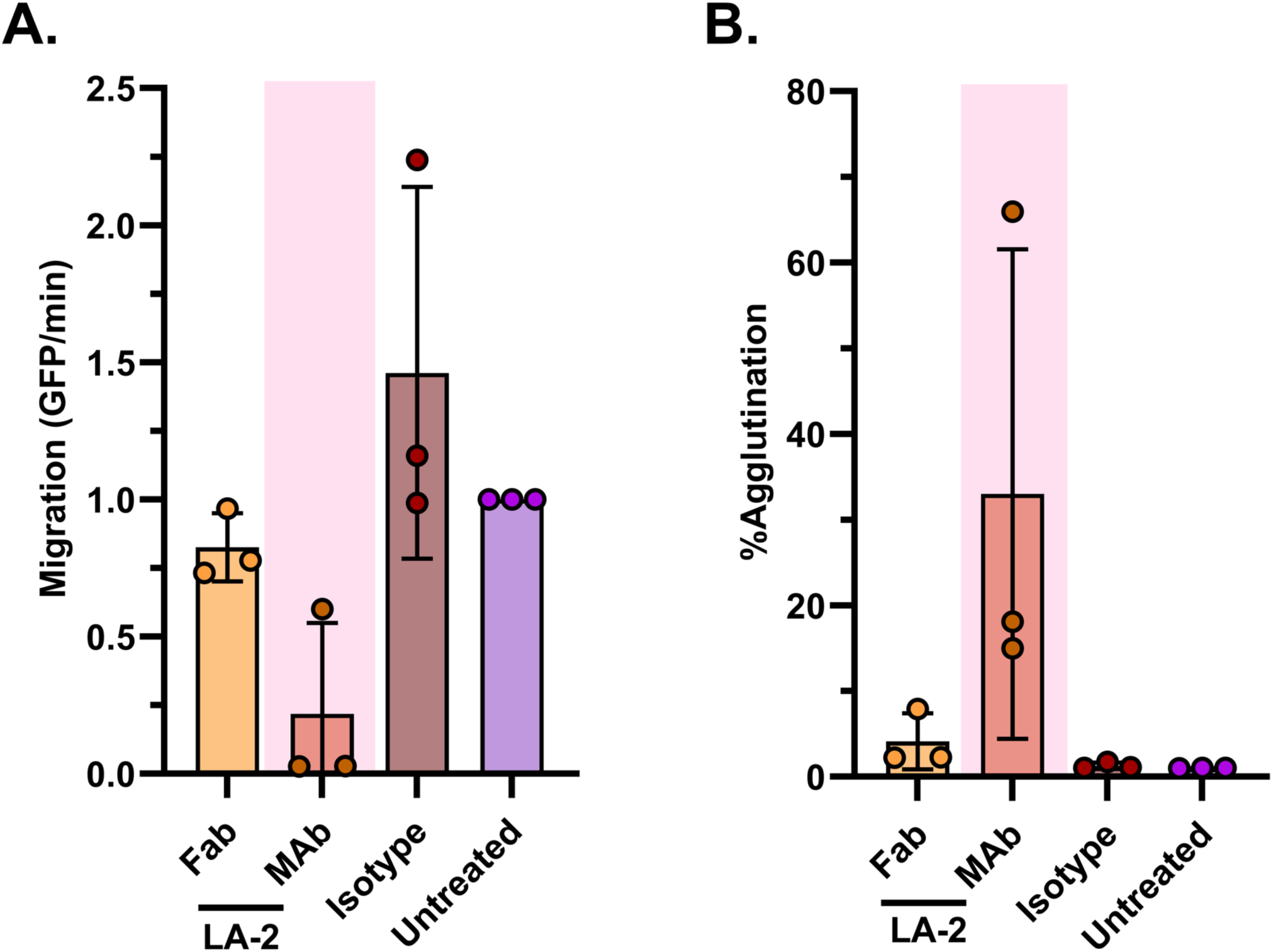
LA-2 monovalent Fab’ fragments fail to significantly inhibit the migration of *B. burgdorferi* B31-A and agglutinate spirochetes in Transwell assay. 2×10^7^ GFP-tagged spirochetes were either untreated or treated with 10 µg/mL of LA-2 Fab’, mAb, or an isotype control mAb in the Transwell assay (Refer to Materials and Methods). Data were acquired from three independent biological replicates. **(A)** Data represents migration (GFP events/min) normalized to untreated spirochetes and error bars represent standard deviation of the mean. **(B)** Data represent percent agglutination of spirochetes in the lower Transwell chamber normalized to untreated spirochetes and error bars represent standard deviation of the mean. Statistical significance was determined by one-way ANOVA followed by Dunnett’s *post hoc* multiple comparisons test. Pink shading indicates *p*≤0.05 compared to isotype mAb-treated group.

While the transmigration assays with LA-2 were done in the absence of exogenous complement in the BSK-II (upper chamber) or low motility media (lower chamber), we were concerned that trace amounts of complement may have carried over from the *B. burgdorferi* B31-A culture. The presence of even trace amounts of complement might have borreliacidal (or bacteriostatic) effects on spirochetes and thereby impact migration. To address this concern, we repeated the transmigration assays with an “Fc silent” derivative of LA-2 known as LA-2 LALAPG. LA-2 LALAPG carries three point mutations in the Fc region of IgG1 that render the molecule unable to fix complement (Palmer *et al*., 2025). We found that LA-2 LALAPG functioned as effectively as LA-2 IgG in transmigration inhibition (**Figure S2**), thereby demonstrating the effects are independent of any Fc activity and likely restricted to Fv region of the antibody.

### LA-2-induced agglutination of *B. burgdorferi* inversely correlates with loss of transmigration activity

A hallmark of certain OspA antibodies, including LA-2, is their capacity to induce agglutination of *B. burgdorferi* cells in culture (Sears *et al*., 1991, Wilske *et al*., 1992, Sadziene *et al*., 1993, Jiang *et al*., 1994, Luft *et al*., 2002, Frye *et al*., 2022). We reasoned that agglutination might account, at least in part, for the lack of spirochete transmigration observed in the Transwell assays in the presence of LA-2. To examine this possibility, we collected the volumes in the lower chambers of the Transwell units in which *B. burgdorferi* B31-A had been treated with 100 µg mL^-1^ LA-2 IgG or an equivalent concentration of an isotype control. By dark-field microscopy, LA-2 IgG treated cells formed large, dense, multicellular aggregates even as cells on the periphery of the aggregates retained wriggling behavior (**Figure 2C**). In contrast, spirochetes treated with 100 µg mL^-1^ of an IgG1 isotype control appeared as single, free-swimming cells that were indistinguishable from cells in the untreated volume (**Figure 2C**). These observations suggest that agglutination may indeed be a factor in influencing spirochete transmigration. In a separate experiment, we treated spirochetes with 100 µg mL^-1^ LA-2, and observed their behavior immediately after, instead of incubating them according to the Transwell assay conditions. We found one end of the spirochetes looping back on itself, while the other end remained free, consistent with what Frye et al. described as “lariats” (Frye *et al*., 2022).

We employed a microvolume flow cytometry protocol developed in our laboratory to quantitate spirochete agglutination as a function of LA-2 IgG concentration (Frye *et al*., 2022, Rudolph *et al*., 2024). Using this assay, we noted a sizeable shift in the forward scatter (FSC) and side scatter (SSC) values upon the addition of LA-2 consistent with agglutination (**Figure 2B**). We observed that LA-2 at 10 µg mL^-1^ resulted in ∼40% spirochetes agglutination, whereas agglutination was negligible (<5%) at lower LA-2 concentrations (**Figure 3B**). Under the same conditions, LA-2 Fab’ (10 µg mL^-1^) induced ∼2% of the spirochetes to agglutinate (**Figure 4B**).

### Impact of OspA human MAbs *B. burgdorferi* transmigration

We next tested whether other OspA mAbs besides LA-2 affect transmigration of *B. burgdorferi* in the Transwell assay. We previously characterized a panel of human OspA mAbs with varying epitope specificities and binding affinities, including three (319-44, 221-7, 857-2) shown to passively protect mice from tick-mediated *B. burgdorferi* challenge (**Table 1**) (Wang *et al*., 2016, Schiller *et al*., 2021, Frye *et al*., 2022, Haque *et al*., 2022). When tested at 10 µg mL^-1^ in the Transwell assay, 10 of the 12 OspA mAbs significantly reduced *B. burgdorferi* migration, albeit to varying degrees (30-90%). The three mAbs (LA-2, 319-44 and 3-24) that target epitopes in Bin 3, which is located at the C-terminus of OspA, had the most pronounced impact on spirochete migration with inhibition values ranging from 89-93% (**Figure 5A**). The two mAbs in Bin 2, 212-55 and 319-28, which recognize opposite faces of OspA’s central β-sheet (*β*-strands 11-13), reduced spirochete transmigration by ∼65% and ∼35%, respectively. Finally, the Bin 1 MAbs, which target OspA’s central *β*-sheet (*β*-strands 8-11) had the most varied impacts: 227-2 reduced migration by ∼70%, 857-2 by 50% and 221-20 by just 37%. The impact on migration was not related to mAb binding affinity, as 857-2’s K_D_ is 20-fold lower than 227-2’s (0.6 versus 10.6 nM), yet 227-2 was more effective than 857-2 at inhibiting bacterial migration. When tested at 1 µg mL^-1^, none of the mAbs had any significant effect on spirochete migration (**Figure S4**).

**Figure 5.**
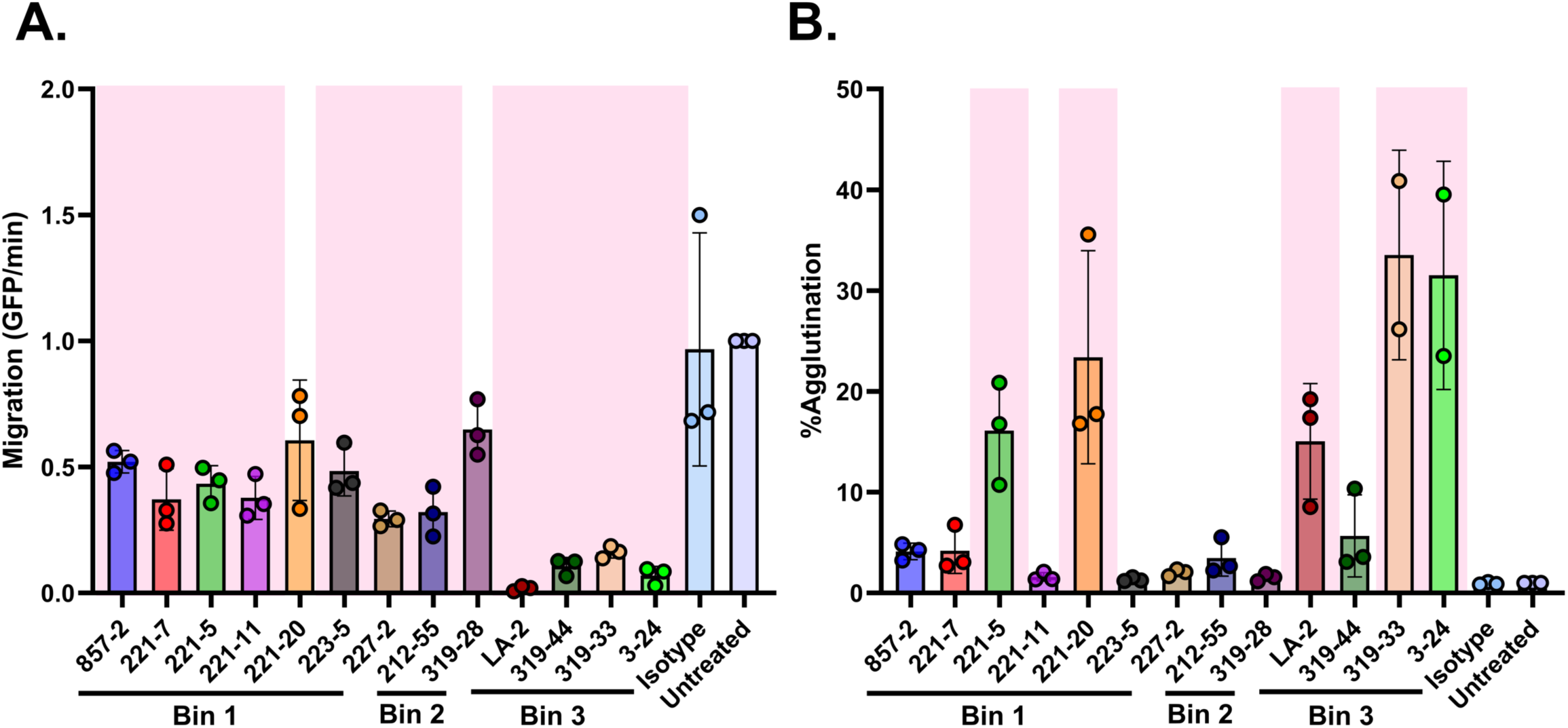
A panel of OspA mAbs inhibit the migration of *B. burgdorferi* B31-A in an agglutination-dependent and - independent manner. 2×10^7^ GFP-tagged spirochetes were either untreated or treated with 10 µg/mL of individual OspA mAbs or an isotype control mAb in the Transwell assay (Refer to Materials and Methods). Data were acquired from three independent biological replicates, excluding 319-33 and 3-24 which were done twice. **(A)** Data represent migration (GFP events/min) normalized to untreated spirochetes and error bars represent standard deviation of the mean. **(B)** Data represent percent agglutination of spirochetes in the lower Transwell chamber normalized to untreated spirochetes and error bars represent standard deviation of the mean. Statistical significance was determined by one-way ANOVA followed by Dunnett’s *post hoc* multiple comparisons test. Pink shading indicates *p*≤0.05 compared to isotype mAb-treated group.

**Figure 6.**
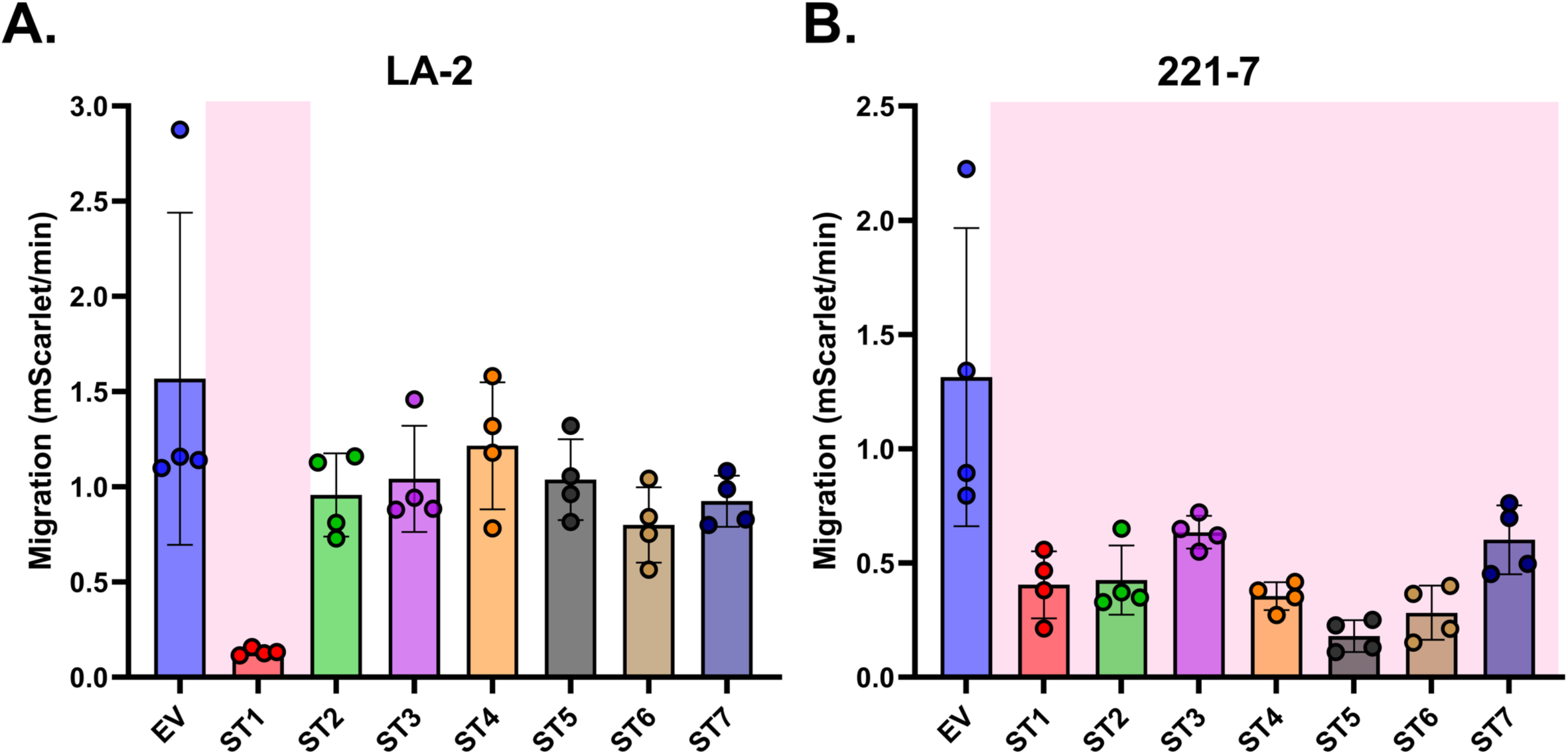
Selected Bin 1 mAbs inhibit the migration of *Borrelia* with different OspA serotypes while LA-2, a Bin 3 mAb, only inhibits migration of serotype 1. 2×10^6^ mScarlet-tagged spirochetes were either untreated or treated with 10 µg/mL of 221-7 or LA-2 in the Transwell lower chamber. The Transwell migration assay was performed as described in Materials and Methods. Data were acquired from three independent biological replicates. Data represent migration (mScarlet events/min) of *B. burgdorferi* HB19-R1 strains (lacks lp54, Δ*ospAB*), with expression plasmids for OspA serotypes 1-7 (ST 1-7), normalized to spirochetes carrying the empty vector (EV), and error bars represent standard deviation of the mean. Statistical significance was determined by one-way ANOVA followed by Dunnett’s *post hoc* multiple comparisons test. Pink shading indicates *p*≤0.05 when compared to empty vector.

To examine the relationship between transmigration inhibition and agglutination in the extended panel of mAbs, the volumes in the lower chamber of the Transwell units were collected at 20 h and subjected to flow cytometry, as described above. A total of five mAbs induced notable spirochete agglutination (**Figure 5B**): three from Bin 3 (LA-2, 3-24 and 319-33) and two from Bin 1 (221-5 and 221-20). These results demonstrate that inhibition of transmigration can occur independent of agglutination, as defined by flow cytometry.

### OspA MAbs inhibit transmigration of *B. burgdorferi* expressing different OspA serotypes

In the United States, Lyme disease is principally associated with *B. burgdorferi* sensu stricto carrying OspA serotype 1 (Wilske *et al*., 1993). However, in Europe and Asia, *Borrelia* genospecies carrying six additional OspA serotypes (ST2-7) are associated with disease (Lee *et al*., 2024). The serotypes are particularly polymorphic towards the C-terminus of OspA corresponding to epitope Bin 3 and more conserved within Bin 1. We therefore tested 221-7, a Bin 1 MAb, along with LA-2 as a control, for its ability to impact the transmigration of *B. burgdorferi* HB19-R1 strains, individually carrying each of the seven OspA serotypes. As expected, LA-2 inhibited the migration of HB19-R1 strain expressing OspA serotype 1 (∼92%) but not the other serotypes. 221-7 at 10 µg mL^-1^ inhibited the migration of all seven serotypes. The strongest effect was seen on the migration of HB19-R1 expressing OspA serotype 5, which represents *B. garinii*.

### Impact of OspA camelid-derived V_H_Hs *B. burgdorferi* transmigration

In addition to the human MAbs described above, we also have a collection of single-domain camelid antibodies (V_H_Hs) that target a diversity of epitopes on OspA and a range of binding affinities as monovalent V_H_Hs or bivalent VHH-IgG1 Fc (Vance *et al*., 2024). We tested three V_H_Hs and their corresponding V_H_H-Fcs each from Bins 1 and 3. At 10 µg mL^-1^, none of the monovalent V_H_Hs had any significant impact on spirochete transmigration (**Figure 7A, C**). Similarly, at 10 µg mL^-1^, none of the VHH-IgG1 Fc had inhibitory activity except for L8H8-IgG1 Fc, which reduced spirochete transmigration by ∼75% (**Figure 7A**). L8H8-IgG1 Fc also promoted spirochete agglutination (∼12%) but at rather low levels (**Figure 7B**). It is probably not a coincidence that L8H8 has the highest binding affinity among all the OspA V_H_Hs we identified.

**Figure 7.**
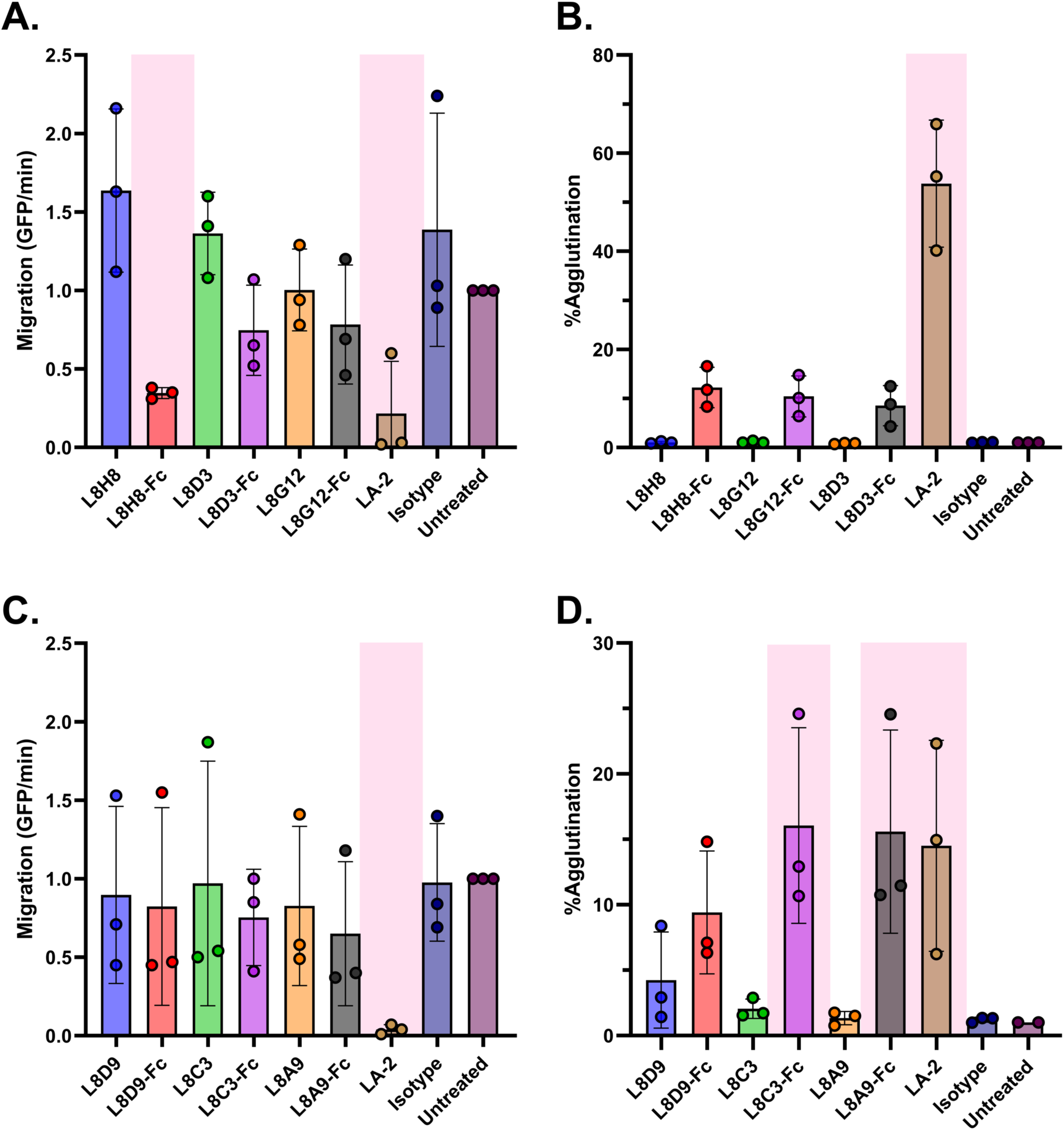
Impact of Bin 1 and Bin 3 representative V_H_H and V_H_H-IgG constructs on the migration of *B. burgdorferi* B31-A and its correlation with spirochete agglutination. 2×10^7^ GFP-tagged spirochetes were either untreated or treated with 10 µg/mL of either V_H_H or V_H_H-IgGs or an isotype control mAb in the Transwell lower chamber. LA-2 at 10 µg/mL was used as the positive control. The Transwell migration assay was performed as described in Materials and Methods. Data were acquired from three independent biological replicates. **(A, C)** Data represent migration (GFP events/min) of spirochetes treated with Bin 1 and Bin 3 V_H_H and V_H_H-IgGs, respectively, normalized to untreated spirochetes, and error bars represent standard deviation of the mean. **(B, D)** Data represent percent agglutination of spirochetes treated with Bin 1 and Bin 3 V_H_H and V_H_H-IgGs, respectively, in the lower Transwell chamber normalized to untreated spirochetes and error bars represent standard deviation of the mean. Statistical significance was determined by one-way ANOVA followed by Dunnett’s *post hoc* multiple comparisons test. Pink shading indicates *p*≤0.05 when compared to isotype mAb-treated group.

### OspA immune sera inhibits *B. burgdorferi* transmigration *in vitro*

In light of the effect of OspA mAbs on *B. burgdorferi* transmigration, we were prompted to evaluate OspA immune sera in the same set-up. Towards this end, we pooled hyper-immune sera from mice that had received several doses of a heptavalent mRNA vaccine formulation (kindly provided by Dr. Meredith Finn, Moderna) which was then diluted 1:100, mixed with *B. burgdorferi* B31-A, and seeded into the lower well of a Transwell unit. The addition of the OspA immune serum resulted in >99% reduction in spirochete migration over an 18 h period relative to cells treated with naïve sera (**Figure 8A**). Similar reduction in migration was seen when it was compared to that of untreated spirochetes. Analysis by dark-field microscopy of the volume in the lower chambers revealed numerous *Borrelia* aggregates of varying sizes, consistent with antibody-induced agglutination (**Figure 8B**). Serial dilutions of the OspA hyperimmune sera (1:200 - 1:2000) revealed a significant inhibition of transmigration at 1:400 dilution, but not beyond that. When the volume in the lower chamber of the Transwell was subjected to flow cytometry, we observed significant macro-agglutination of spirochetes at the 1:200 and 1:400 dilutions, but not at lower doses of the serum (**Figure 8C**). We conclude that OspA polyclonal sera inhibits *B. burgdorferi* transmigration *in vitro* at concentrations associated with spirochete agglutination.

**Figure 8.**
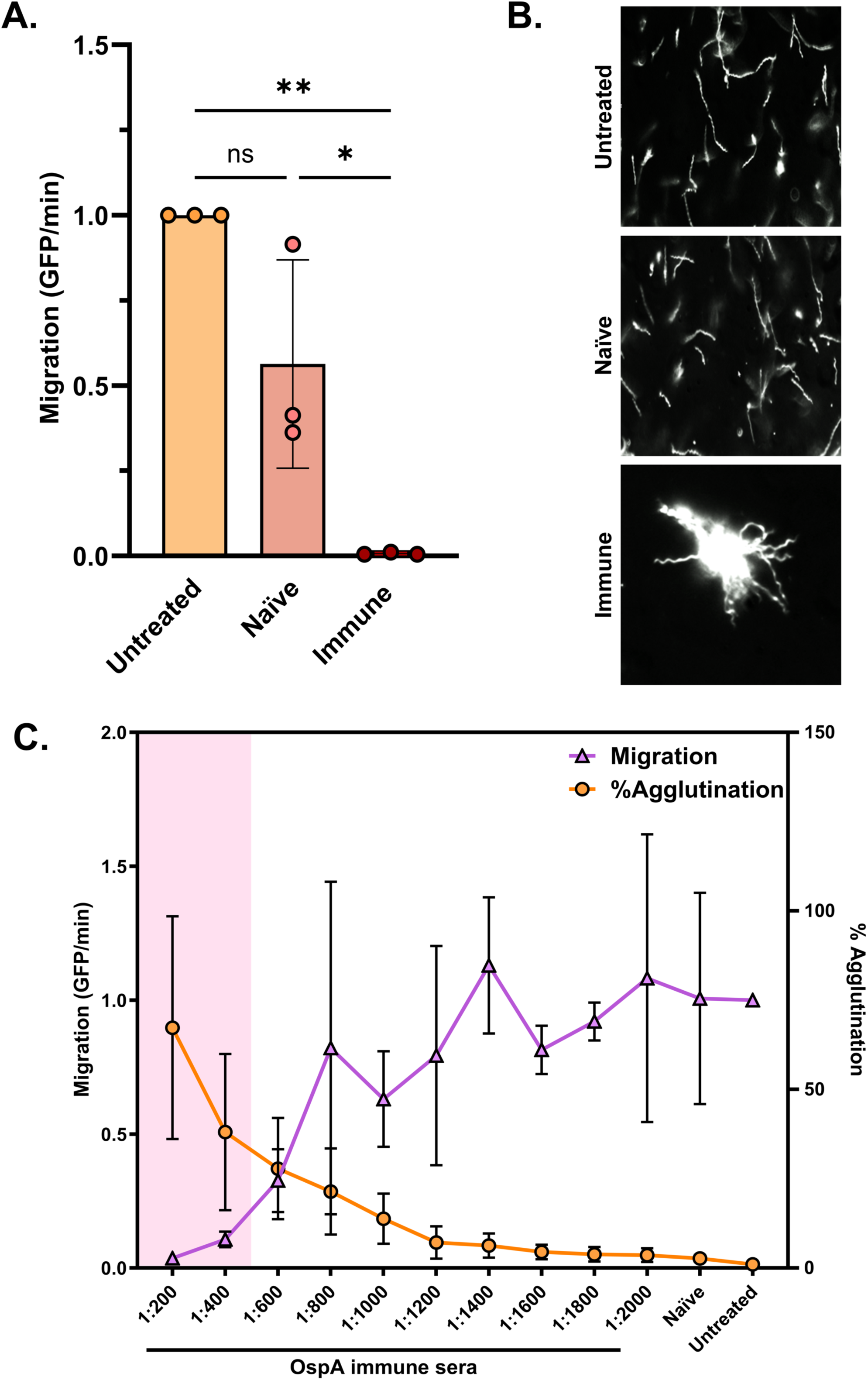
Polyclonal sera from immunized mice significantly inhibits the migration of *B. burgdorferi* B31-A, in a dose-dependent manner which correlates with antibody-mediated spirochete agglutination. (A) 2×10^7^ GFP-tagged spirochetes were either untreated or treated with 1:100 dilutions of naïve or immune serum. The Transwell migration assay was performed as described in Materials and Methods. Data were acquired from three independent biological replicates. Data represent migration (GFP events/min) normalized to untreated spirochetes and error bars represent standard deviation of the mean. Statistical significance was determined by one-way ANOVA followed by Tukey’s post hoc multiple comparisons test. Asterisks indicate *p*≤0.05 (*) and *p*≤0.01 (**) and ns=not significant. **(B)** Dark field microscopy images of spirochetes in the Transwell lower chamber after overnight incubation with 1:100 serum dilution. All images have been captured at 20X objective magnification. **(C)** 2×10^7^ GFP-tagged spirochetes were either untreated or treated with a range of immunized serum dilutions or 1:200 dilution of naïve serum. The Transwell migration assay was performed as described in Materials and Methods. Data were acquired from three independent biological replicates. Data represent migration (GFP events/min, in violet) and percent agglutination (in orange), normalized to untreated spirochetes and error bars represent standard deviation of the mean. Statistical significance was determined by one-way ANOVA followed by Dunnett’s post hoc multiple comparisons test. Pink shading indicates *p*≤0.05 when compared to naïve serum-treated group.

## DISCUSSION

The observation that OspA antibodies entrap *B. burgdorferi* within the tick midgut during the course of a bloodmeal was one of several pivotal findings that contributed to the eventual development of OspA-based Lyme disease vaccines (Fikrig *et al*., 1990, de Silva *et al*., 1996, Steere *et al*., 1998, Gomes-Solecki *et al*., 2020, Plotkin & Shapiro, 2024). Subsequent reports demonstrated that transmission blocking activity of OspA antibodies does not require host complement or impact spirochete viability (Rathinavelu *et al*., 2003, Gipson & de Silva, 2005). Since those seminal studies were conducted more than 20 years ago, few if any reports have sought to address how it is that OspA antibodies interfere with *B. burgdorferi* egress from the tick midgut without impacting spirochete viability.

In this study, we investigated the effect of OspA monoclonal and polyclonal antibodies on *B. burgdorferi* migration across a porous (3-micron) polycarbonate membrane in a Transwell unit, as a proxy for spirochete movement out of the tick midgut. The transmigration assay we employed is a modification of a method developed by Van Gundy and colleagues for the purpose of identifying *B. burgdorferi* chemoattractants in tick salivary glands (Van Gundy *et al*., 2021). Using this set-up, we demonstrated that LA-2 IgG is a potent inhibitor of *B. burgdorferi* transmigration when seeded into the lower chamber along with the spirochetes. LA-2 is arguably one of the most well-characterized transmission-blocking OspA mAbs in the field (Schaible *et al*., 1990, Ding *et al*., 2000, Wang *et al*., 2016, Shandilya *et al*., 2017, Frye *et al*., 2022, Haque *et al*., 2022). LA-2 targets the C-terminus of OspA, including the exposed loops between β-strands 16-17, 18-19, and 20-21 (Ding *et al*., 2000). From the standpoint of Lyme disease vaccines, LA-2 is highly significant as LA-2 equivalents, based on competitive ELISAs, correlate with protection from *B. burgdorferi* challenge in animal models (Johnson *et al*., 1995, Golde *et al*., 1997). An LA-2 equivalence assay was subsequently employed in human clinical trials of the original OspA-based vaccine (Steere *et al*., 1998).

While the exact physiological relevance of our results remains to be determined, there is considerable evidence that the concentrations of LA-2 that inhibited spirochete transmigration in the Transwell unit are within the range associated with transmission inhibition *in vivo*. For example, the “LA-2-like” mAb, C3.78 IgG, passively protected mice from tick-mediated *B. burgdorferi* infection at serum antibody concentrations roughly corresponding to 10 µg mL^-1^ (Gipson & de Silva, 2005). In case of humans vaccinated with lipidated full-length OspA, individuals with serum LA-2 equivalent titers of 10 µg eq mL^-1^ or higher had no incidence of Lyme disease (Steere *et al*., 1998). However, in the case of both monoclonal and polyclonal antibodies, defining an absolute threshold associated with protection is difficult considering that numerous extrinsic factors influence transmission, including spirochete numbers in the tick midgut at the time of antibody exposure (Golde *et al*., 1997) and levels of tick proteins in the midgut known to impact migration and transmission efficacy (Coumou *et al*., 2016).

Examination of a well-characterized collection of human mAbs targeting different epitopes along the length of OspA revealed that virtually all of them could inhibit (to varying degrees) *B. burgdorferi* movement across the Transwell membrane. As a rule, mAbs that targeted epitopes overlapping with LA-2 (so called Bin 3) were more potent at inhibiting transmigration than mAbs that recognize epitopes elsewhere on the molecule (Bins 1 and 2). The Bin 3 mAbs were also the most effective at promoting spirochete agglutination, as determined by flow cytometry and dark field microscopy. This agrees with previous observations from our group in which we demonstrated that LA-2 was significantly more effective than 857-2 (Bin 1 mAb) at inducing *B. burgdorferi* agglutination in culture (Frye *et al*., 2022). We speculate that large (macroscopic) bacterial aggregates are not conducive to passage through 3-micron pores in the Transwell membrane. However, in that same report, it was noted that agglutination is likely the culmination of a series of antibody-induced changes in *B. burgdorferi* behavior that initiate with the formation of “lariats,” in which one pole of a spirochete loops back onto itself forming a loop. While lariat formation does not affect flagella rotation per se, as evidenced by the fact that the spirochetes continue to wriggle under these conditions, the bacteria do seemingly lose their directionality and displacement (Harman *et al*., 2012, Frye *et al*., 2022). It has been reported that antibodies drive OspA to localize at the spirochete poles, possibly because of alterations in membrane fluidity and localization of cholesterol rafts (Toledo *et al*., 2014). While the interrelationships between OspA mis-localization, lariat formation, and *B. burgdorferi* motility are yet to be determined, it is tempting to speculate that they contribute to changes in transmigration.

We recognize that the Transwell unit does not recapitulate transmigration of *B. burgdorferi* across the tick midgut epithelium *in vivo*. As detailed in a recent review, during the course of a blood meal, the tick midgut is lined by a chitin-rich membrane that acts as a physical barrier to limit particle and pathogen access to epithelial cells (Kitsou *et al*., 2021). In the Transwell unit, the spirochetes presumably move unimpeded through the porous membrane. Moreover, spirochetes residing in the midgut transition to a motile, invasive phenotype during tick feeding (Dunham-Ems *et al*., 2009). In the Transwell model, the lower chamber is seeded with “fully energized” *B. burgdorferi* cells collected from mid-log phase cultures grown in BSK II. Finally, antibody exposure in the tick midgut is presumably highly variable and subject to fluctuations in volume during a feeding event. This contrasts with the Transwell unit where *B. burgdorferi* cells are effectively bathed in a uniformly high concentration of OspA-specific antibodies prior to initiating transmigration. Nonetheless, the observation that both monoclonal and polyclonal OspA antibodies can arrest *B. burgdorferi* movement across a physical barrier in a dose and time dependent manner independent of complement or other blood components is consistent with mechanisms proposed by others to explain transmission inhibition *in vivo* (Rathinavelu *et al*., 2003, Gipson & de Silva, 2005).

## Acknowledgements

We are grateful to Drs. Taylor Van Gundy and Meghan Lybecker at the Centers for Disease Control and Prevention (Fort Collins, CO) for assistance in establishing the transmigration assay. We thank Dr. Graham Willsey (Wadsworth Center) for providing *B. burgdorferi* strains, Carol Lyn Piazza (Wadsworth Center) for assistance with flow cytometry protocols, and Dr. David Vance (Wadsworth Center) for providing VHHs and guidance on antibody selection. We thank Dr. Meredith Finn (Moderna, Inc) for sharing OspA immune mouse sera. We gratefully acknowledge Drs. Renjie Song and Jennifer Yates of the Wadsworth Center’s Immunology Core for assistance with flow cytometry and the Media and Cell Culture core for BSK-II medium and *Borrelia* motility buffer. We thank Ms. Elizabeth Cavosie (Wadsworth Center) for administrative assistance. The authors have no conflicts of interest to declare.

