## Supplemental figures for "OspA Antibodies Inhibit the In vitro Transmigration of *Borreliella burgdorferi*"

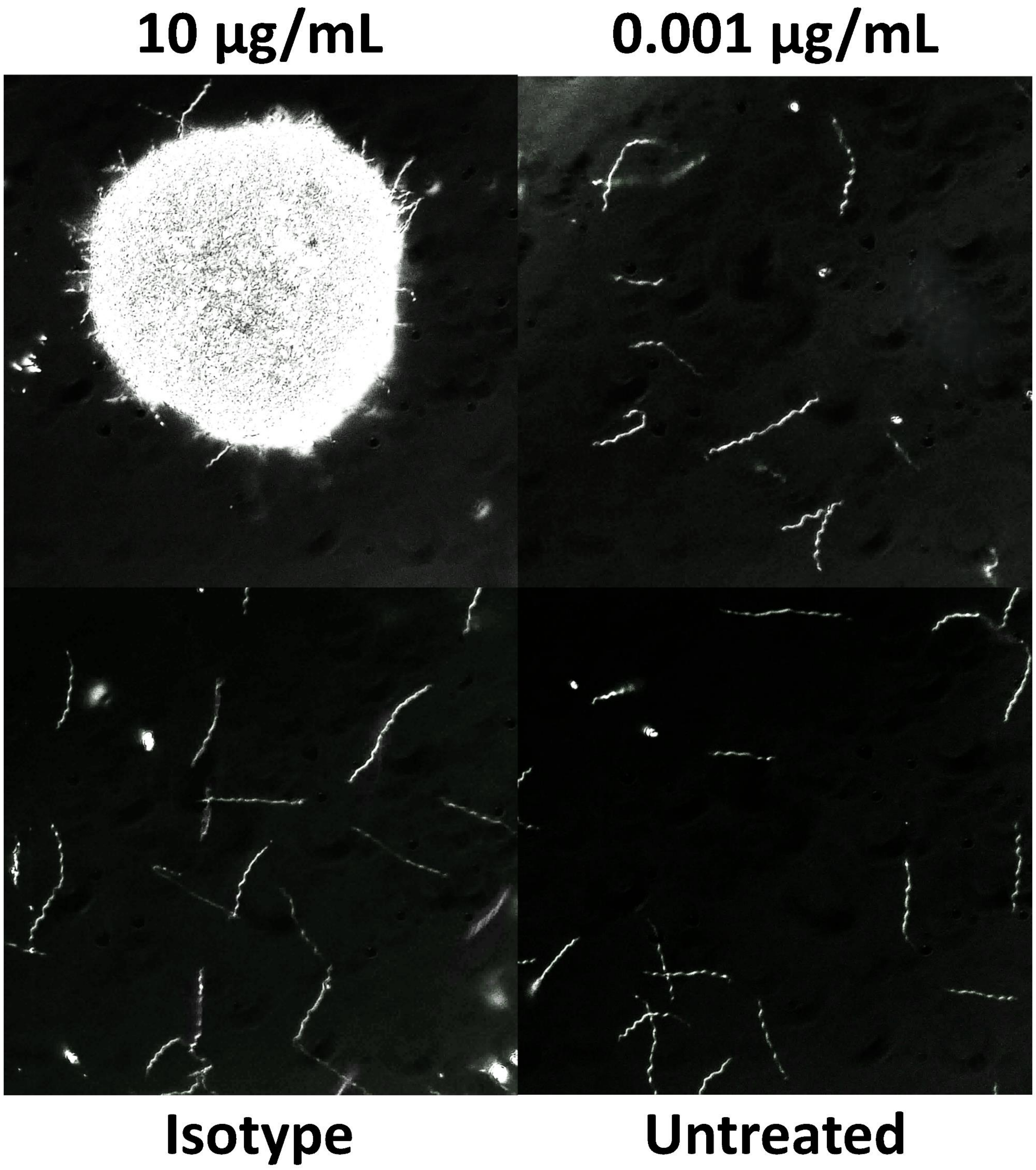


**Figure S1. Dark field microscopy images of *B. burgdorferi* B31-A exposed to varying concentrations of LA-2.** Representative images show spirochetes treated with the highest and lowest concentration of LA-2 in the dose response range, or in control conditions in the Transwell assay. All images have been captured at 20X objective magnification.

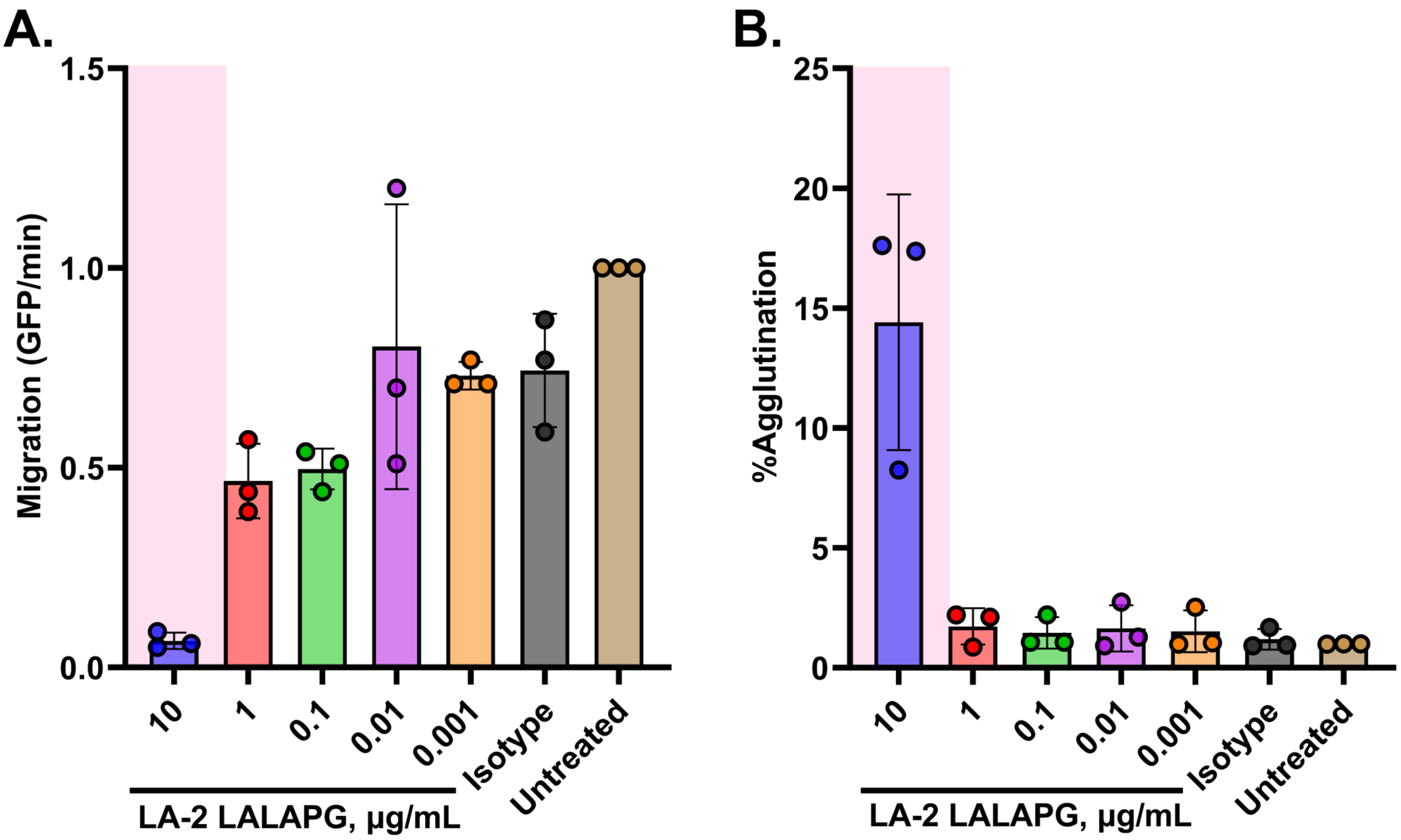


**Figure S2. A Fc-silent variant of LA-2 (LA-2 LALAPG) inhibits the migration of *B. burgdorferi* B31-A in a dose-dependent manner.** 2×10$^{7}$ GFP-tagged spirochetes were either untreated or treated with a range of LA-2 LALAPG concentrations or 10 µg/mL of isotype control mAb in the Transwell lower chamber. The Transwell migration assay was performed as described in Materials and Methods. Data was acquired from three independent biological replicates. **(A)** Data represents migration (GFP events/min) normalized to untreated spirochetes and error bars represent standard deviation of the mean. (B) Data represents percent agglutination of spirochetes in the lower Transwell chamber normalized to untreated spirochetes and error bars represent standard deviation of the mean. Statistical significance was determined by one-way ANOVA followed by Dunnett's *post hoc* multiple comparisons test. Pink shading indicates $p \leq 0.05$ and compared to isotype mAb-treated group.

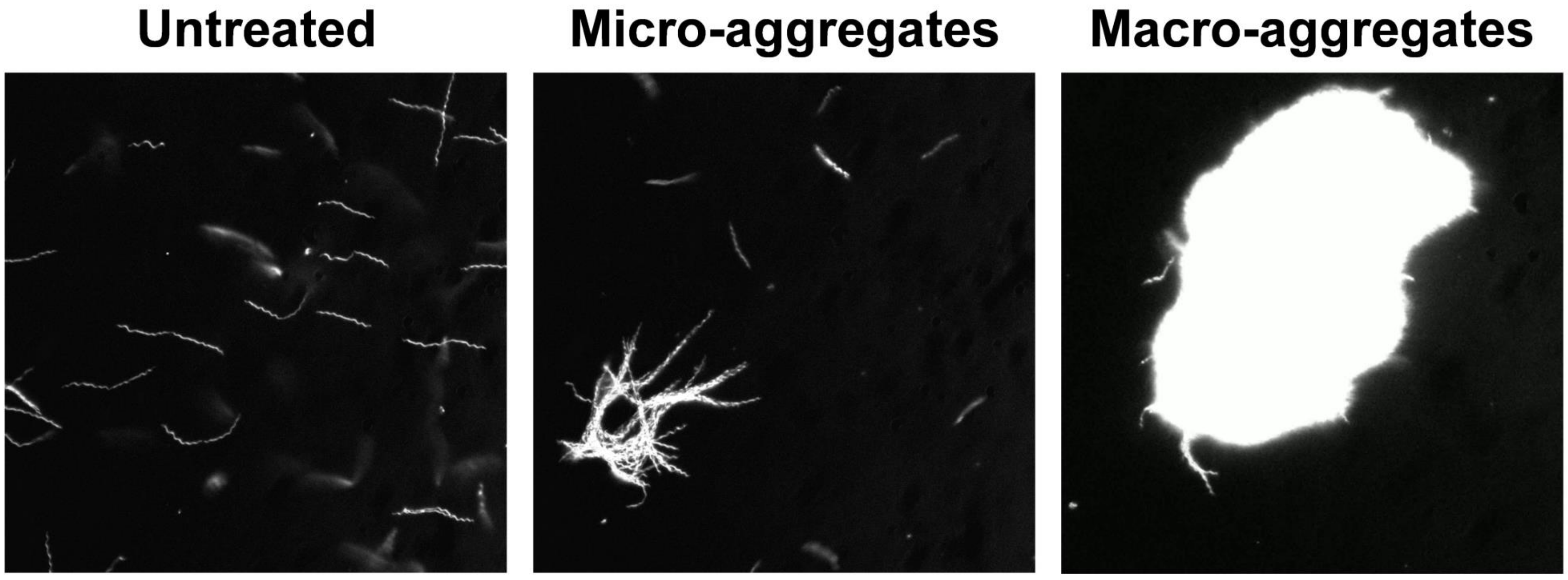


**Figure S3. Dark field microscopy images of *B. burgdorferi* B31-A exposed to different OspA mAbs.** Representative images show untreated spirochetes and the formation of micro-aggregates, mostly by Bin 1 and Bin 2 mAbs, and macro-aggregates, mostly by Bin 3 mAbs, with a few exceptions. All images have been captured at 20X objective magnification.

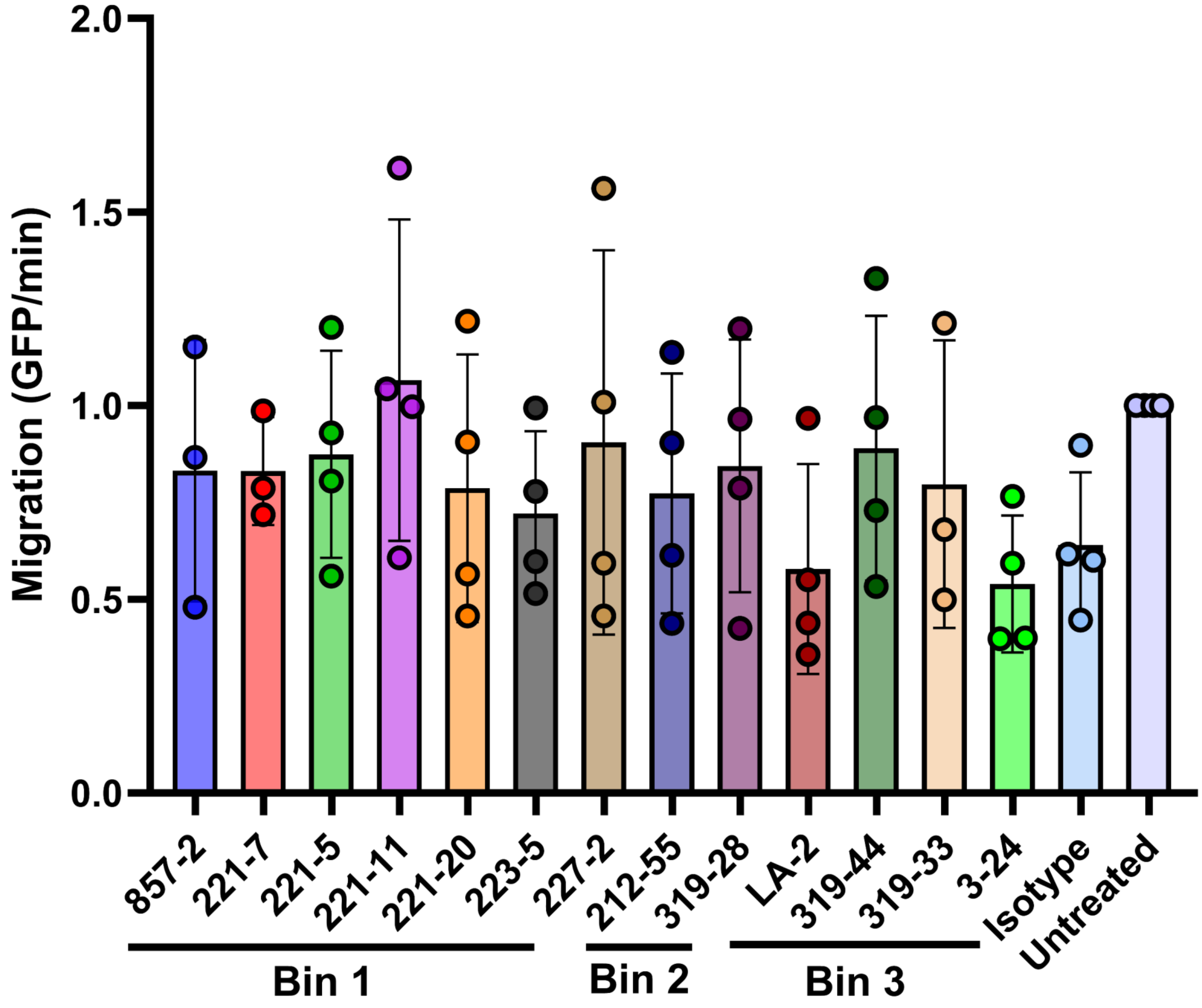


**Figure S4. A panel of OspA mAbs fail to inhibit the migration of *B. burgdorferi* B31-A at a low dose.** $2{\times}10^7$ GFP-tagged spirochetes were either untreated or treated with 1 µg/mL of individual OspA mAbs or an isotype control mAb in the Transwell assay (Refer to Materials and Methods). Data was acquired from three independent biological replicates. Data represents migration (GFP events/min) normalized to untreated spirochetes and error bars represent standard deviation of the mean. There was no statistically significant difference compared to isotype mAb-treated group, as determined by one-way ANOVA followed by Dunnett's *post hoc* multiple comparisons test.
